# SSUplex: fast, both-strand extraction and origin-sorting of small-subunit rRNA for environmental DNA metabarcoding

**DOI:** 10.64898/2026.07.02.736232

**Authors:** Aaron O’Brien, Juan Vargas Botia, Isabel Acuña, Franko Restovic, Patricio Martínez, Pilar Parada

## Abstract

Ribosomal RNA metabarcoding sits at the center of how we characterize microbial and eukaryotic communities in environmental samples, and long-read sequencing has made full-length small-subunit (SSU; 16S/18S) profiling routine. The broadly conserved primers that make rRNA such a convenient marker are also its liability: by design they co-amplify organellar (mitochondrial, chloroplast) and cross-domain SSU alongside the intended target. Left unsorted before taxonomic assignment, these passengers are systematically misclassified, and the error propagates straight into estimates of community composition and diversity. Reads must therefore be detected, extracted, and sorted by origin before they ever reach a classifier. We present SSUplex, an open-source tool that detects SSU rRNA, assigns each read to one of five origins (bacteria, archaea, eukaryota, mitochondria, chloroplast), and extracts the SSU region for downstream classification. SSUplex reimplements the extraction-and-origin logic of the widely used Metaxa2 in the Rust programming language, scans both strands, and ships as a single dependency-light binary suited to long-read (Oxford Nanopore, PacBio HiFi) and short-read data. Benchmarked against Metaxa2 on public data, SSUplex reproduces Metaxa2 origin calls on full-length reads (96.8% concordance) and matches its extraction speed on small inputs, then pulls away to run up to ∼3.4× faster with ∼35% lower peak memory at 200,000 reads, the per-sample scale a long-read amplicon run typically reaches. We are candid about a genuine, measured trade-off in the origin-ranking statistic, and we pinpoint the bacteria-versus-mitochondria boundary as the method’s one intrinsically lower-confidence edge. For the now-common workflow in which origin-sorted reads are handed to a dedicated classifier rather than classified in place, SSUplex is a fast, reproducible, embeddable stand-in for Metaxa2’s extraction role. Source code and a benchmark harness that regenerates every result from public data are available under the MIT license at https://github.com/ayobi/ssuplex.

## 1 Introduction

Metabarcoding of ribosomal RNA marker genes is a workhorse of community ecology and microbiology, used to survey bacterial and archaeal assemblages (16S), microbial eukaryotes and fungi (18S), and to track how these communities shift as their environment does. As long-read sequencing has matured, it has carried the approach up to full-length SSU genes, buying taxonomic resolution that the short hypervariable windows of short-read platforms simply cannot reach.

Between the raw reads and any biological inference, though, sits a practical obstacle. The “universal” primers we rely on to capture rRNA markers are, by design, broadly conserved, and that same conservation lets them pull in far more than the intended target: eukaryotic 18S in a nominally bacterial 16S run and, because the organelle ribosome traces its ancestry to bacteria, the SSU genes of mitochondria and chloroplasts. Plant-associated, host-associated, and many soil samples are especially thick with these organellar reads. Hand them straight to a bacterial reference classifier and chloroplast reads come back as *Cyanobacteria*, mitochondrial reads as *Rickettsiales*, and eukaryotic reads as low-confidence noise. What results is no cosmetic annotation slip but a distortion of the very quantities a study sets out to measure: relative abundances, richness, and diversity. Sorting reads by biological origin before they reach a classifier is, for this reason, a standard and necessary preprocessing step.

Metaxa2 [1] has been the reference implementation of this step since 2015. It detects SSU and LSU genes using hidden Markov models (HMMs) of their conserved subregions, reads origin from the pattern of region matches, and then runs BLAST-based taxonomic classification. The same conserved-flank strategy drives ITSx [6] for the internal transcribed spacer (ITS); indeed, SSUplex stands to Metaxa2’s SSU extraction much as our earlier Rust tool ITSxRust [7] stands to ITSx for ITS, and together they cover the principal rRNA metabarcoding markers. Two properties, however, hold Metaxa2 back on contemporary datasets. First, it is Perl orchestration wrapped around HMMER and BLAST, last released in 2021; the BLAST classification stage, which dominates runtime, fires on every invocation even when the user fully intends to classify the extracted reads with a different, marker-specific tool. Second, a single long-read run can now return millions of full-length SSU reads, and in that regime per-read overhead becomes the practical bottleneck for the extraction step itself.

Here we describe SSUplex, a focused reimplementation of the SSU extraction-and-origin step in Rust, engineered around the read counts and read orientations of long-read environmental data (Figure 1). SSUplex does detection and origin assignment and nothing more, leaving taxonomic classification to whichever dedicated tool a study has chosen.

Several other tools detect rRNA in sequence data but occupy different niches. RNAmmer [3] and barrnap [4] predict rRNA gene loci in assembled genomes and metagenomes (the latter, like SSUplex, built on HMMER profiles), but they are oriented toward annotating contigs rather than sorting individual reads by domain of origin. SortMeRNA [5] separates rRNA reads from non-rRNA in metatranscriptomic data, which is a different decision (rRNA versus messenger RNA) from the cross-domain origin assignment we need here. Metaxa2 remains the tool aimed squarely at detecting the SSU region in environmental reads and resolving its bacterial, archaeal, eukaryotic, mitochondrial, or chloroplast origin, and it is therefore the natural baseline for SSUplex. Table 1 lays out how these tools differ in task, input, and output.

## 2 Implementation

### Detection

SSUplex accepts FASTA input, optionally gzip-compressed. SSU rRNA is detected with profile HMMs of the conserved subregions of the gene. These are assembled from the Metaxa2 SSU profile database into one set per origin: the bacterial, archaeal, eukaryotic, and chloroplast profiles are taken directly, while Metaxa2’s separate metazoan-mitochondrial (*mitozoa*) profiles are folded together with the mitochondrial set so that all mitochondrial signal is scored as one origin. Each read is searched against every profile set with `nhmmer` from the HMMER 3 suite [2], which scans both the forward and reverse-complement strands. Hits are parsed from the `nhmmer` tabular output; envelope coordinates are normalised so that minus-strand hits are expressed on forward-strand coordinates, and hits weaker than an E-value cutoff (default 10^*−*5^, set with `--evalue`) are discarded. Both-strand search is not optional for long-read data, where orientation is effectively a coin flip: in development, restricting the search to the forward strand recovered only 2,622 of 5,000 mock-community reads (52%), whereas scanning both strands

**Table 1:**
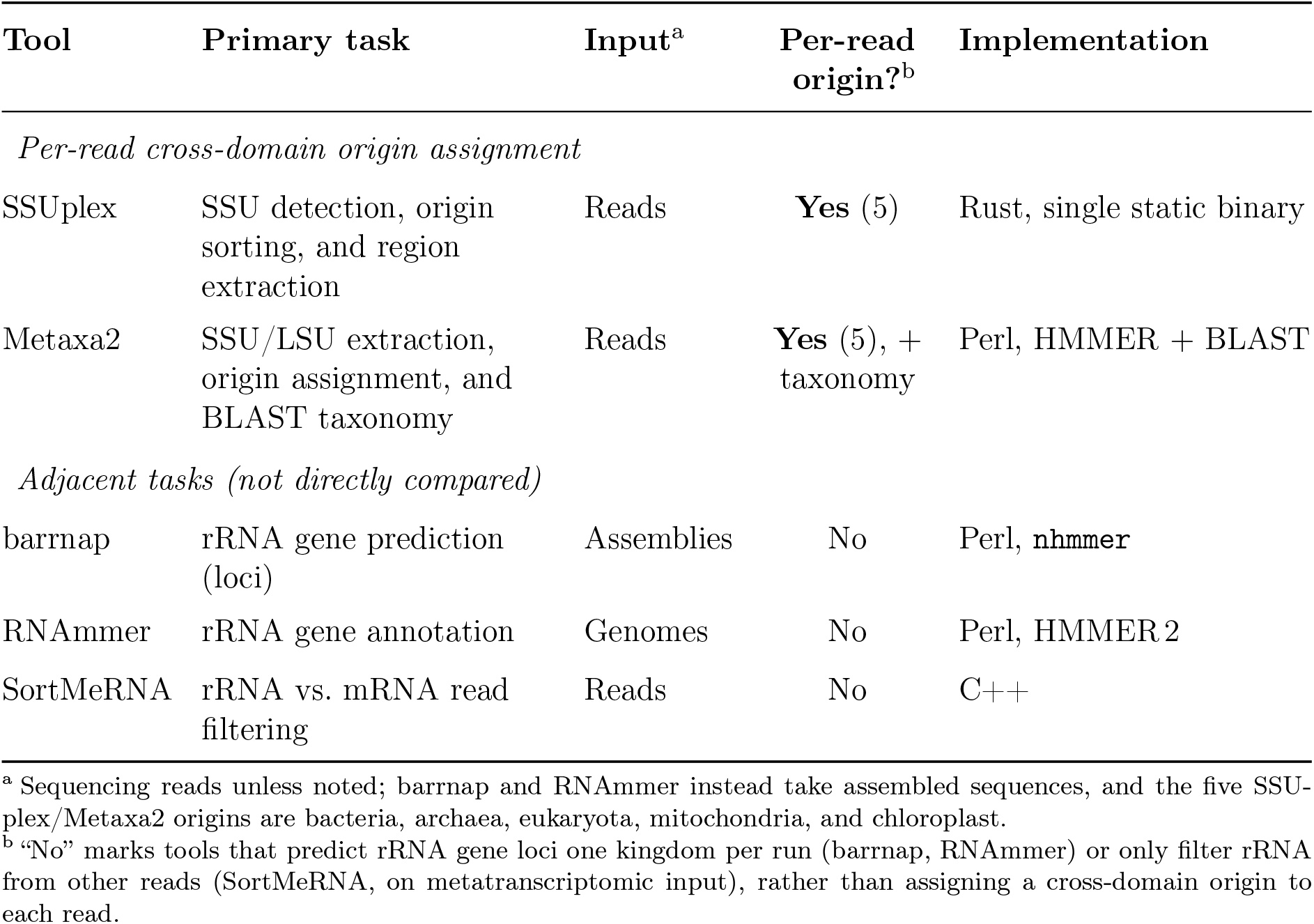
Scope of SSUplex relative to other rRNA sequence tools. Only SSUplex and Metaxa2 assign a cross-domain origin to individual SSU reads and so share SSUplex’s task; the remaining tools solve adjacent but distinct problems, which is why the quantitative comparison in this paper is made against Metaxa2 alone.

### Origin assignment

For a read *r* and origin *o*, the surviving hits define *n*_*r,o*_, the number of matched conserved-region profiles, with summed bit score *S*_*r,o*_ and mean per-region bit score 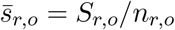. SSUplex assigns the origin

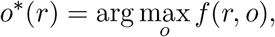

where the ranking statistic *f* is the mean 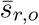 (default), the sum *S*_*r,o*_, or the count *n*_*r,o*_, selected with `--rank`. The assignment is accepted only if 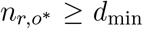 (`--min-domains`, default 1) and 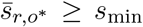 (`--min-score`, default 0); failing either test, the read is reported as unclassified rather than forced into an origin it does not earn. Ties are resolved by the number of matched regions and then by a fixed ordering of origin labels, so assignments are fully deterministic.

### Extraction and output

Accepted reads are clipped to the SSU coordinate envelope defined by their HMM hits and written to one FASTA file per origin, ready to hand to a downstream classifier. SSUplex also writes a per-read table of origin call, region count, score, and coordinates, and a run summary; a diagnostic mode emits the full per-read, per-origin score table and the raw per-region hits.

### Engineering and availability

SSUplex is implemented in Rust and distributed as a single statically linked binary with no language runtime dependency; its only external requirement is `nhmmer` on the `PATH`. Profile search is parallelised across CPU threads and input is streamed. The tool builds from a single Rust source tree and has been compiled and run end-to-end on both x86-64 Linux and Intel macOS; as an indication of its modest requirements, on a four-core Intel MacBook (16 GB RAM) it extracted SSU from 200,000 full-length ONT reads in roughly 13 min on four threads, using about 1.3 GB of memory. Source code, documentation, and the benchmark harness are available under the MIT license at https://github.com/ayobi/ssuplex.

**Figure 1:**
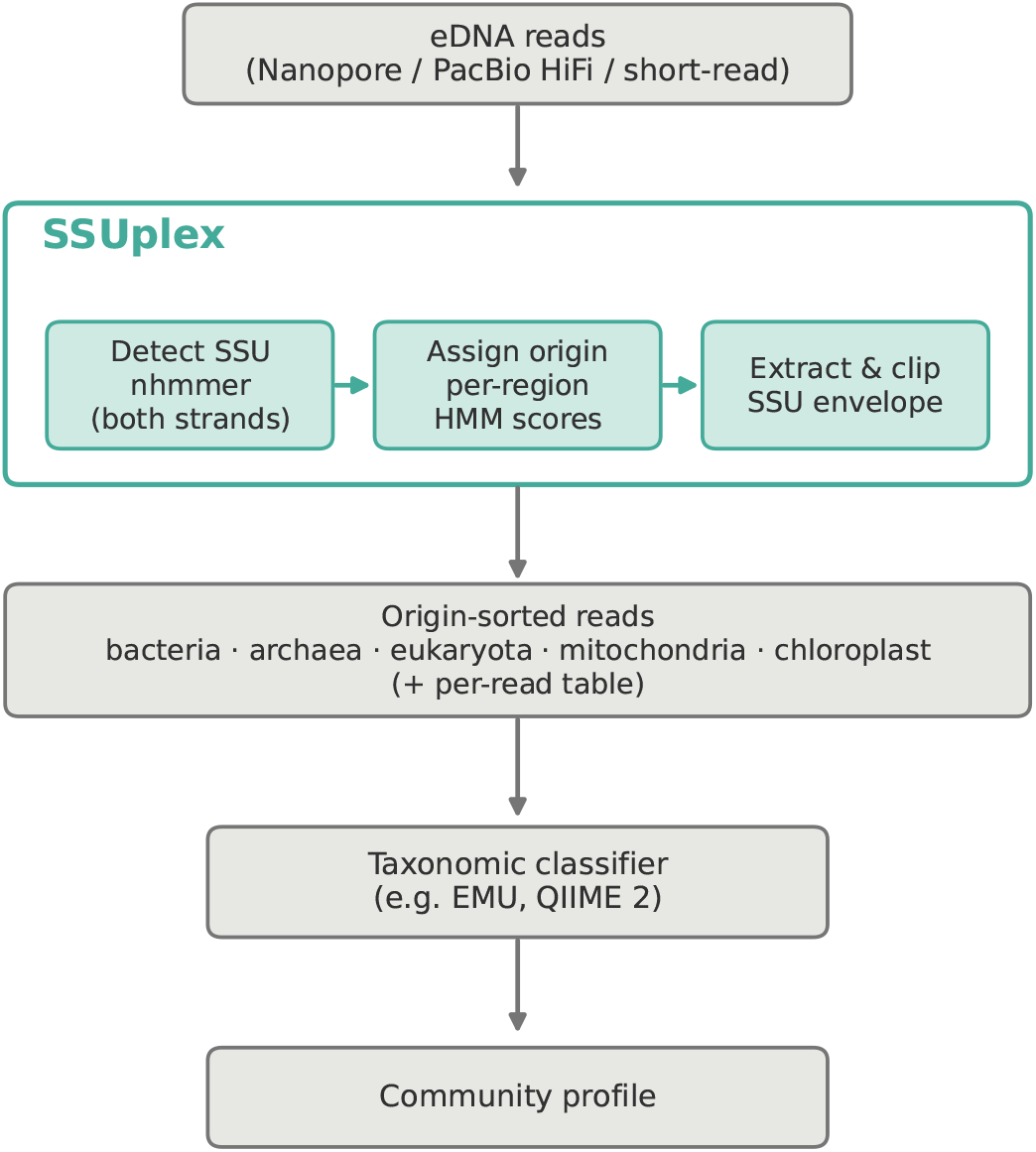
SSUplex in an rRNA-marker workflow. Reads are scanned on both strands to detect SSU rRNA, assigned to one of five origins from their per-region HMM scores, and clipped to the SSU envelope. The origin-sorted reads (and a per-read table) are handed to a dedicated taxonomic classifier; SSUplex performs no classification itself.

## 3 Results

We evaluated SSUplex against Metaxa2 (v2.2.3) with a reproducible harness distributed with the source; Metaxa2 serves throughout as the reference standard, and the harness regenerates every input from public data.

### 3.1 Origin-assignment concordance

We assessed origin calls on three datasets spanning clean and realistic conditions: a full-length SSU reference set sampled from the Metaxa2 profile database (clean reads, known origin); the ZymoBIOMICS Oxford Nanopore mock community (SRA run SRR10391201; all-bacterial ground truth); and a real plant-root Nanopore 16S sample (rice endophytes; BioProject PRJNA992961, run SRR25243163), an organelle-rich, host-associated sample for which no per-read truth exists and we therefore report concordance with Metaxa2.

The ranking statistic buys us a genuine, measured trade-off (Table 2). On clean full-length reads all three statistics perform well and the mean is marginally best. On the bacteria-only mock, the summed score is far more accurate than the mean, because on noisy reads a spurious organelle origin can post a higher *mean* over a few regions while the true bacterial origin matches *more* regions at a slightly lower mean. On the organelle-rich plant sample the picture flips: the mean tracks Metaxa2 closely on organellar reads, agreeing on 1,302 of 1,741 reads Metaxa2 called mitochondrial and 1,871 of 1,987 it called chloroplast, whereas the summed and count statistics misroute abundant plant-mitochondrial reads to bacteria. No single statistic wins across every regime. We ship the mean as the default, because it most faithfully reproduces Metaxa2 on the multi-origin samples typical of environmental data, and we expose `--rank sum` for bacteria-dominated samples such as gut or many soil communities.

**Table 2:** Origin-assignment performance by ranking statistic (--rank), showing the trade-off that no single statistic resolves. The two numeric columns are accuracy against known ground truth; the rice column reports qualitative concordance with Metaxa2, for which no per-read truth exists.

| Statistic | Accuracy vs. ground truth |  | Concordance |
| --- | --- | --- | --- |
|  | Reference<sup>a</sup> | ZymoBIOMICS<sup>b</sup> | Rice, multi-origin<sup>c</sup> |
| mean (default) | **96.8%** | 43.0%<sup>d</sup> | Tracks Metaxa2 on mitochondrial and chloroplast reads |
| sum | 95.6% | **95.9%** | Misroutes abundant plant-mitochondrial reads to bacteria |
| count | 78.0% | 95.7% | Over-collapses organellar reads to bacteria |

<sup>a</sup> Clean full-length reads sampled from the Metaxa2 SSU profile database (known origin).
<sup>b</sup> ZymoBIOMICS ONT mock community; all-bacterial ground truth.
<sup>c</sup> Rice plant-root ONT 16S sample; organelle-rich, no per-read truth, so agreement with Metaxa2 is reported rather than accuracy.
<sup>d</sup> Not a failure of detection: on this bacteria-only, noisy set a spurious organelle can win a higher *mean* over a few regions, whereas the true bacterial origin matches *more* regions at a slightly lower mean, which is exactly why `sum` is preferred for bacteria-dominated samples.

### 3.2 Application to a plant-root sample

To show the triage task on exactly the kind of data that motivates it, we ran SSUplex with default settings on the 5,000-read rice plant-root sample. It assigned 1,141 reads (22.8%) to bacteria, 1,602 (32.0%) to mitochondria, 2,248 (45.0%) to chloroplast, 7 to archaea, and 2 to eukaryota. Roughly three-quarters of the reads in this nominally bacterial 16S sample are, then, organellar; sent straight to a bacterial reference classifier they would have been misassigned, whereas SSUplex peels them off into dedicated files for appropriate (or no) downstream classification. This breakdown is what one expects of the organelle-rich composition of plant-associated material and, as above, reflects concordance with Metaxa2 rather than independent ground truth.

### 3.3 Extraction throughput and memory

We compared extraction throughput on the ZymoBIOMICS Nanopore reads across a 40× range of input sizes (Figure 2). Both tools ran in extraction-only mode (SSUplex by default; Metaxa2 with `-x T`), both scanning both strands, both with 12 threads, on a single workstation; the smallest point was run in triplicate (under 2% variance). At small inputs the two are neck and neck, but the gap opens with scale, reaching ∼3.4 at 200,000 reads and still widening, while SSUplex draws ∼35% less memory and occupies fewer CPU cores (∼4 versus ∼6.5). Extrapolating the observed, approximately linear scaling (dashed lines, Figure 2) puts this ratio near ∼3.5× at 500,000–1,000,000 reads, with the absolute time difference continuing to grow; we present these as projections to convey scale, not as measurements. That range spans the persample read counts of long-read amplicon studies. We stress that this compares the extraction step only; Metaxa2’s full pipeline additionally runs BLAST classification, which SSUplex omits by design and which accounts for the far larger end-to-end differences seen when Metaxa2 runs in its default mode.

**Figure 2:**
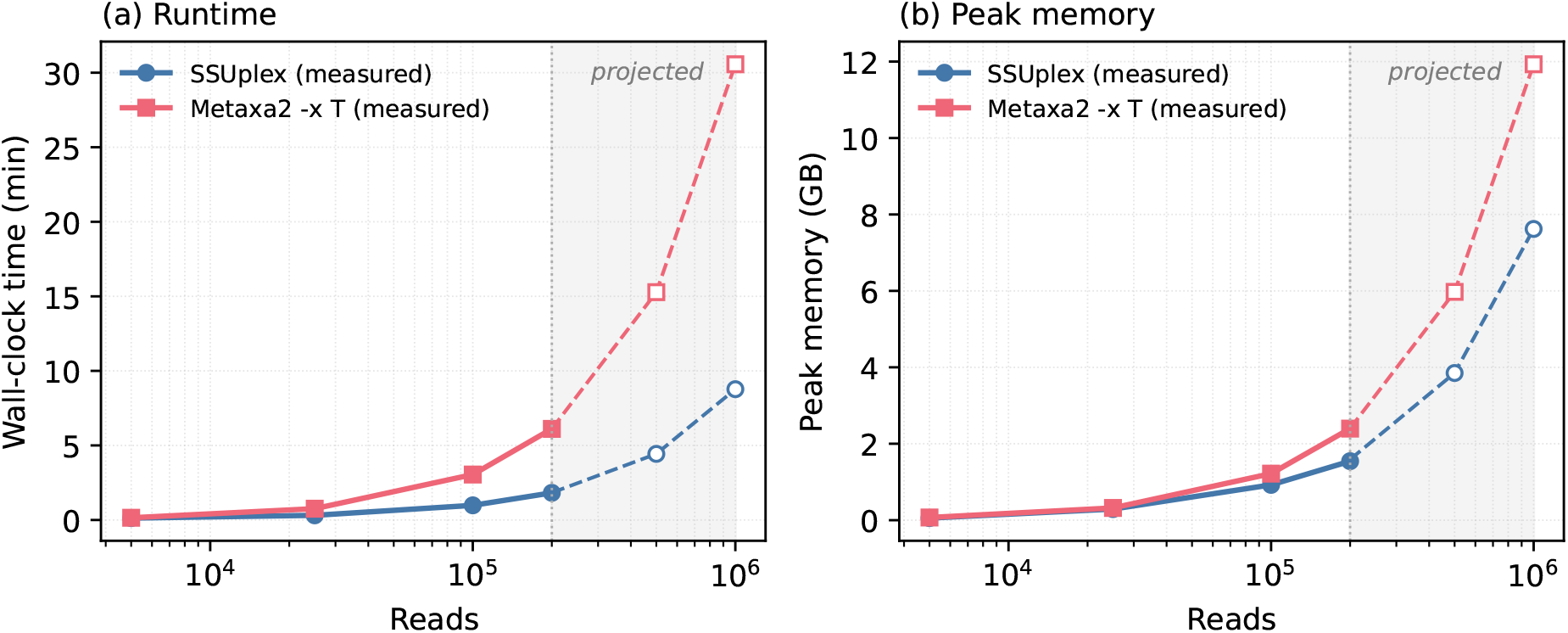
Extraction-only performance versus input size on ZymoBIOMICS Nanopore reads (12 threads, single workstation; Metaxa2 run with `-x T`, both tools scanning both strands). (a) Wall-clock runtime (minutes) and (b) peak memory. Filled markers and solid lines are measured values up to 200,000 reads; open markers and dashed lines in the shaded region are a linear extrapolation to 500,000 and 1,000,000 reads, shown to indicate scale and *not measured here*. The measured runtime gap reaches ∼3.4× at 200,000 reads; assuming the observed approximately linear scaling continues, the gap holds near ∼3.5× while projected peak memory approaches ∼8 GB (SSUplex) and ∼12 GB (Metaxa2) at one million reads. SSUplex performs no taxonomic classification.

Multithreading is implemented across the five origin profiles, so it delivers a ∼3–4× speedup and saturates near four threads (28.5 s at one thread, 9.2 s at four, 8.2 s at twelve, on 5,000 reads); asking for more threads buys nothing further. Peak memory scales with read count because sequences are indexed in memory, which is comfortable at amplicon scale but becomes the limiting factor for multi-million-read shotgun metagenomes (projected peak memory ~8 GB for SSUplex and ∼12 GB for Metaxa2 at one million reads; Figure 2b), and it is precisely this that motivates the streamed-input work noted in the Discussion.

### 3.4 Scope and limitations

The bacteria-versus-mitochondria call is the method’s one intrinsically lower-confidence edge. Because mitochondrial SSU is of alpha-proteobacterial origin, the two are genuinely close on noisy reads, and per-read inspection shows that the winner can hinge on how many weak, near-threshold conserved regions survive, a decision no single HMM-score statistic resolves cleanly. The per-read score margins make this plain (Figure 3): reads Metaxa2 attributes to chloroplast or mitochondria carry a positive organelle-minus-prokaryote mean-score margin and bacterial reads a negative one, but the distributions bleed into each other near zero. Under SSUplex’s own scores, 34% of reads Metaxa2 calls bacterial and 13% of those it calls mitochondrial land on the opposite side of the decision boundary, precisely the reads a single statistic cannot resolve. Metaxa2 disambiguates them with its second, BLAST-based classification stage, which SSUplex deliberately leaves out. Mitochondrial-versus-bacterial calls on noisy data should therefore be read as lower-confidence, and the default mean statistic is the one to prefer when mitochondrial signal matters. Where SSUplex is strong, by contrast, it is strong and fast: at SSU detection and extraction on both strands, at telling chloroplast, archaeal, and eukaryotic SSU apart across clean and noisy data, and at bacterial assignment on clean reads and on bacteria-dominated samples with `--rank sum`.

**Figure 3:**
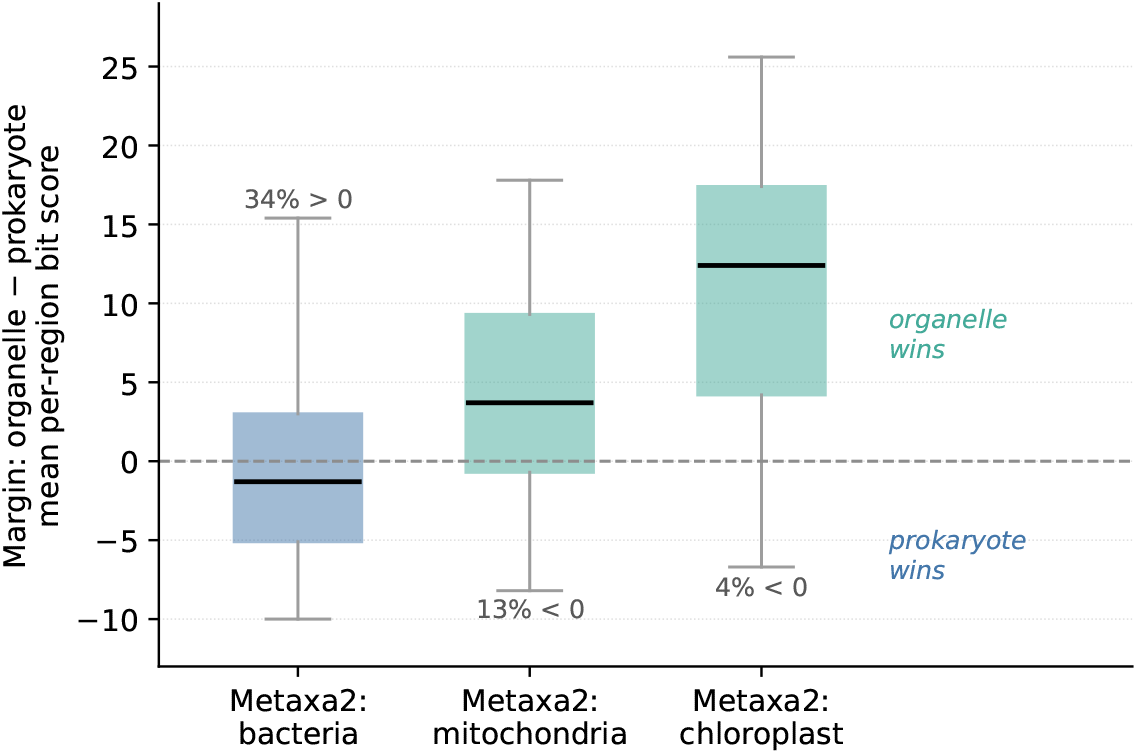
Origin-score margin separability on the plant-root sample, using SSUplex’s own perread scores grouped by the origin Metaxa2 assigned. The margin is the best organelle minus the best prokaryote mean per-region bit score; a positive margin selects an organellar origin under the default mean ranking. Box spans the 10th–90th percentile, line is the median, whiskers the range. Chloroplast and mitochondrial reads separate from bacterial reads across the zero boundary, but overlap near it, exactly the bacteria/mitochondria cases that no single HMM-score statistic resolves.

### 3.5 Benchmark details and reproducibility

Benchmarks used Metaxa2 v2.2.3, HMMER 3 [2], and SSUplex v0.1.0 on a single workstation (AMD Ryzen 9 9950X, 16 cores / 32 threads; 64 GB RAM; Ubuntu). The origin HMM profile sets were assembled from the Metaxa2 SSU database with the script provided in the repository. Extraction was run identically for both tools, scanning both strands at 12 threads:

~~~
ssuplex -i reads.fasta -o out/s --hmm-dir hmms -t 12
metaxa2 -i reads.fasta -o out/run -x T --cpu 12 --multi_thread T --silent T
~~~

Runtime and peak memory were recorded with `/usr/bin/time -v`; the 5,000-read point was measured in triplicate (variance < 2%) and reported as the median. The reference truth set was generated from the Metaxa2 SSU database with the harness’s read simulator, with optional fragmentation and sequencing-error injection; the ZymoBIOMICS (SRR10391201) and rice (SRR25243163) reads were streamed directly from the European Nucleotide Archive. Origin concordance was computed by joining the per-read origin calls of the two tools and tabulating agreement. All inputs, commands, and analysis scripts are included in the repository so that every table and figure here can be regenerated.

## 4 Discussion

SSUplex is best understood not as a wholesale replacement for Metaxa2 but as a scoped one. For the now-common workflow in which extracted, origin-sorted reads go to a dedicated long-read classifier rather than being classified in place, SSUplex steps directly into Metaxa2’s extraction role: it reproduces Metaxa2 origin assignment on full-length reads, applies the same both-strand handling, and delivers substantially higher throughput at lower memory at the read volumes long-read datasets now reach. It does not replace Metaxa2’s integrated taxonomic classification, which it omits on purpose in favour of a clean hand-off, and it does not match Metaxa2 on the bacteria/mitochondria boundary on noisy reads. Within that clearly bounded scope, though, its advantages are concrete and measured: a single static binary in place of a Perl pipeline, faster extraction at realistic scale, lower memory, and a design that never forces a BLAST classification a study may not want.

These properties make SSUplex easy to embed in existing long-read metabarcoding pipelines and to run reproducibly at the per-sample scale of modern surveys, handing origin-sorted reads to a marker-appropriate classifier and so trimming back the organelle- and cross-domain-driven biases that would otherwise propagate into community and diversity estimates. Because it ships as a single dependency-light binary and holds only hundreds of megabytes to a few gigabytes of memory at typical amplicon read counts, it runs happily on commodity laptops and modest servers, with no compute cluster and none of the Perl and BLAST stack Metaxa2 requires, which lowers the barrier for smaller and resource-limited laboratories taking on eDNA studies.

The most useful directions for future work fall straight out of the analysis above. A lightweight mapping-based cross-check for reads near the bacteria/organelle boundary, effectively a fast analogue of Metaxa2’s second stage applied only to the ambiguous reads, would shore up the one place where HMM region scores fall short. An updated archaeal profile panel would improve recovery of under-represented lineages, streamed input would lift the memory ceiling for very large metagenomes, and extension to LSU (23S/28S) extraction would broaden marker coverage. Together with ITSxRust [7] for ITS extraction and EMITS [8] for ITS classification, SSUplex rounds out a coherent, reproducible, open-source toolkit for rRNA-marker analysis of environmental DNA.

## Author contributions

A.O. conceived, implemented, and benchmarked the software and wrote the manuscript. J.V.B., I.A., F.R., and P.M. contributed to the conception of the work through extensive discussions on environmental DNA analysis. P.P. provided direction and supervision as principal investigator. All authors read and approved the final manuscript.

## Data and code availability

SSUplex is available at https://github.com/ayobi/ssuplex under the MIT license, with a benchmark harness that regenerates all inputs used here. No new sequencing data were generated. Sequencing data are public: the ZymoBIOMICS ONT mock community (SRA run SRR10391201; BioProject PRJNA587452) and the plant-root ONT 16S sample (SRA run SRR25243163; BioProject PRJNA992961).

## Acknowledgements

During software development and manuscript preparation, we used Claude (Anthropic) for assistance with code review, writing, and editing, and GitHub Copilot (Microsoft) for code completion and suggestion. All AI-assisted output was critically reviewed and verified by the authors, who take full responsibility for the content of the software and this manuscript.

